# Resistance at No Cost: The Transmissibility and Potential for Disease Progression of Drug-Resistant *M. Tuberculosis*

**DOI:** 10.1101/475764

**Authors:** Mercedes C. Becerra, Chuan-Chin Huang, Leonid Lecca, Jaime Bayona, Carmen Contreras, Roger Calderon, Rosa Yataco, Jerome Galea, Zibiao Zhang, Sidney Atwood, Ted Cohen, Carole D. Mitnick, Paul Farmer, Megan Murray

## Abstract

**Background:** The future trajectory of drug resistant tuberculosis strongly depends on the fitness costs of drug resistance mutations. Here, we measured the association of phenotypic drug resistance and the risk of TB infection and disease among household contacts (HHCs) of patients with pulmonary TB.

**Methods:** We evaluated 12767 HHCs of patients with drug sensitive and resistant pulmonary TB at baseline, two, six, and 12 months to ascertain infection status and to determine whether they developed tuberculosis disease. We also assessed the impact of drug resistance phenotype on the likelihood that a TB strain shared a genetic fingerprint with at least one other TB patient in the cohort.

**Findings:** Among 3339 TB patients for whom were DST available, 1274 (38%) had TB that was resistant to at least one drug and 478 (14⋅3%) had multi-drug resistant (MDR) TB, i.e. TB resistant to both INH and rifampicin. Compared to HHCs of drug sensitive TB patients, those exposed to a patient with MDR-TB had an 8% (95% CI: 4-13%) higher risk of infection by the end of follow up. We found no statistically significant difference in the relative hazard of incident TB disease among HHCs exposed to MDR-TB compared to DS-TB (Adjusted HR 1⋅28 [(95% CI: ⋅9-1⋅83]). Patients with MDR-TB were more likely to be part of a genetic cluster than were DS-TB patients.

**Interpretation:** Clinical strains of MDR M. tuberculosis are neither less transmissible than drug sensitive strains nor less likely to cause disease. (ClinicalTrials.gov number, NCT00676754)

**Funding:** National Institutes of Health: NIH/NIAID CETR U19AI109755

**Statement:** All authors have seen the manuscript and approved the manuscript.

---

Resistance to tuberculosis (TB) drugs threatens to slow gains against the global TB pandemic. In 2015, of 10⋅4 million new TB cases, 500,000 were caused by MDR-TB, i.e., TB resistant to both isoniazid and rifampicin. Access to effective treatment for MDR-TB remains limited, and cure is achieved in only 50%.^1^

The primary strategy deployed against MDR-TB was designed to reduce the emergence of new resistance but did not address existing drug resistant strains. This policy focused on the empirical treatment of presumed drug-sensitive TB (DS-TB) to reduce the acquisition of drug-resistance in individual patients during suboptimal therapy.^2,3^ While this approach likely prevents patients from acquiring new drug resistance, it does nothing to interrupt the transmission of existing DR-TB strains.^4^ With MDR-TB prevalence in some regions as high as 32%,^1^ the course of the epidemic will depend on how quickly DR-TB patients are detected and rendered non-infectious as well as on the relative transmissibility of DR-TB strains.^5–7^ If the acquisition of drug-resistance mutations by Mycobacterium tuberculosis (Mtb) incurs a “fitness cost” that reduces its ability to spread, the incidence of DR-TB would be expected to decline more slowly than if resistance were cost-free.^7–9^ The projections of mathematical models forecasting TB incidence strongly depend on this putative cost.^7^

We undertook to estimate the fitness cost of drug resistance by comparing the rates of TB infection and disease among individuals exposed to DR-TB and DS-TB. We further measured the association between drug resistance profile (DRP) and whether strains were members of genetic clusters.

## METHODS

### Study population and setting

This study was conducted in a catchment area of Lima, Peru that included approximately 33 million residents. Detailed descriptions of recruitment and enrollment procedures are provided in the Supplement. The study protocol was approved by the Harvard University Institutional Review Board and by the Research Ethics Committee of the National Institute of Health of Peru.

#### Index patient assessment

Briefly, we identified adults >=15 years who presented with clinically suspected pulmonary TB at 106 participating health centers from September 2009 through August 2012. Two sputum samples were evaluated by Ziehl-Neelsen (AFB) and mycobacterial culture.

At enrollment, we collected information on age, gender, socio-demographics, BCG vaccination, alcohol and tobacco use, comorbid disease, previous TB disease, duration of symptoms, presence of cavitary disease on chest radiograph, sputum smear status, culture results, HIV status, and CD4 count. For those with positive TB cultures, isolates underwent drug-susceptibility testing (DST) and MIRU-based genotyping.

Because DST results are not routinely available until two or more months after patients start therapy, those with unsuspected DR-TB received ineffective regimens prior to the receipt of DST results and would thus be expected to remain infectious during that period. We therefore also recorded the time from enrollment until the initiation of effective therapy (See Supplement).

#### Enrollment of household contacts

Within two weeks of enrolling an index patient, we invited his or her household contacts (HHCs) to participate in this longitudinal study. We screened consenting HHCs for TB signs and symptoms; those who screened positive were referred to a health center for routine evaluation and treatment.

#### Assessment of household contacts

At enrollment, we obtained the following information: age, gender, socio-demographics, BCG vaccination, alcohol and tobacco use, comorbid disease, concurrent medication use, TB disease history, use of isoniazid preventive therapy (IPT), height, weight, HIV status, and CD4 count. In HHCs with no history of TB, we obtained a tuberculin skin test (TST). We considered HHCs to be TB infected if they had a previous history of TB or a TST induration size >10 mm in non-HIV infected contacts and >5 mm in HIV-infected contacts.

We evaluated all HHCs for pulmonary and extra-pulmonary TB disease at two, six, and 12 months after enrollment. HHCs diagnosed with TB disease were classified as having either co-prevalent TB (disease diagnosed within 14 days of the index patient) or secondary TB (disease diagnosed after 14 days). We defined TB disease in HHCs <18 years according to consensus guidelines for diagnosis of TB in children.^10^ HHCs who were previously TST negative were retested at six and 12 months. We considered HHCs to have become infected during follow-up if they had a negative TST at any point in the study that was followed by an increase in TST induration size of at least 6 mm.

### Statistical Analysis

#### TB transmission

To determine whether DRP of the index patient was an independent risk factor for TB infection among HHCs, we used modified Poisson regressions to assess the prevalence odds of infection by the end of follow-up, thereby capturing infection events that occurred both prior to and after the diagnosis of the index patient. In sensitivity analyses, we assessed the association between DRP and TST positivity at enrollment, and used Cox frailty proportional hazards models to assess the association between DRP and the risk of becoming infected during follow up. We estimated the time of infection as the mid-point between the last negative TST and either the first positive TST or a diagnosis of TB disease. We considered the possibility that the impact of DRP was mediated through its effect on either the clinical presentation of TB in the index patient (sputum smear positivity or cavitary lesions) or through prolonged infectiousness due to a delay in receipt of effective treatment. To assess the direct effect of DRP on infection, i.e. the effect that is not mediated through these intermediates, we compared models with and without adjustment for these factors. Because we considered that children who were infected at baseline were more likely to have been infected by the corresponding index patient rather than during a prior exposure, we also conducted sensitivity analyses that restricted the population of HHCs to children <15.

#### TB disease incidence

To assess the impact of DRP on the risk of secondary TB disease in HHCs, we used a Cox frailty proportional hazards model, adjusting for factors known or hypothesized to affect TB disease risk. We also estimated this effect with and without adjustment of the possible mediators cited above. We conducted sensitivity analyses that included only secondary cases who shared an identical or similar MIRU pattern with the index patient.

#### TB clustering

We assessed the impact of DRP on the genetic clustering of TB cases, under the assumption that clustering is a proxy for transmission and disease progression. We classified *Mtb* isolates as clustered if they shared a 24-locus MIRU pattern with an isolate from at least one other case in the study. We then used logistic regression to estimate the impact of DRP on propensity to cluster.

### Role of the funding source

The funder of the study had no role in study design, data collection, data analysis, data interpretation, or writing of the report. The corresponding author had full access to all study data and had final responsibility for the decision to submit for publication.

## RESULTS

Between September 2009 and September 2012, we enrolled 4,500 index patients, of whom 4,044 had microbiologically confirmed TB disease (Table S1). Of the 3,339 for whom DSTs were available, 1274 (38%) had isolates resistant to at least one drug: 538 (16⋅4%) to only one drug and 478 (14⋅3%) to both isoniazid and rifampicin (MDR-TB). The proportion of patients who had MDR-TB decreased with age, ranging from 34⋅1% among those 0-15 years of age to 6⋅2% in those over 60. We enrolled 10,160 household contacts of the 2,563 microbiologically confirmed index patients who had a DTS available and at least one HHC (Table 1).

**Table 1.**
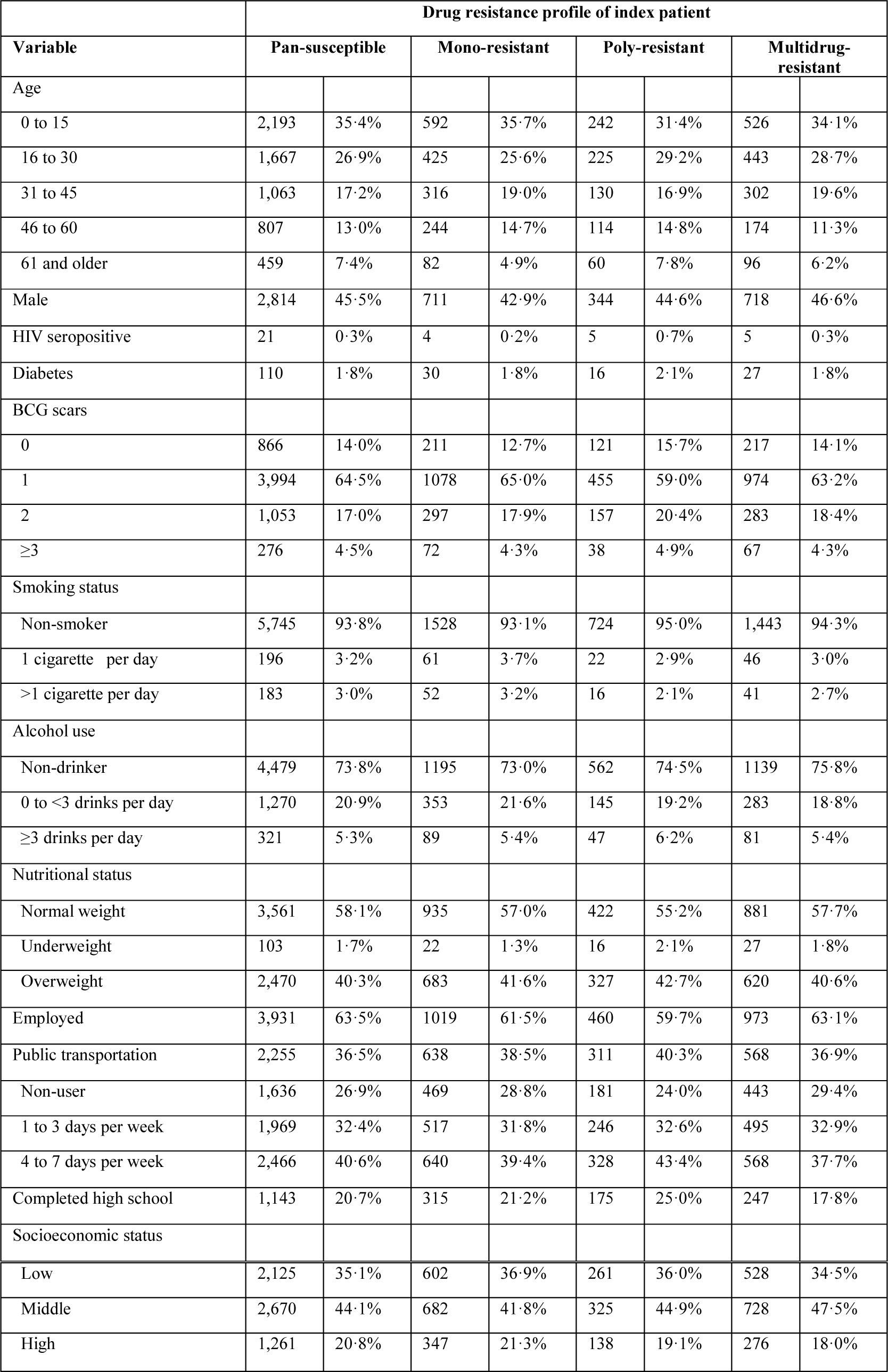
Characteristics of household contacts of patients with pulmonary TB in Lima, Peru.

| Variable | Drug resistance profile of index patient |  |  |  |  |  |  |  |
| --- | --- | --- | --- | --- | --- | --- | --- | --- |
|  | Pan-susceptible |  | Mono-resistant |  | Poly-resistant |  | Multidrug-resistant |  |
| Age |  |  |  |  |  |  |  |  |
| 0 to 15 | 2,193 | 35.4% | 592 | 35.7% | 242 | 31.4% | 526 | 34.1% |
| 16 to 30 | 1,667 | 26.9% | 425 | 25.6% | 225 | 29.2% | 443 | 28.7% |
| 31 to 45 | 1,063 | 17.2% | 316 | 19.0% | 130 | 16.9% | 302 | 19.6% |
| 46 to 60 | 807 | 13.0% | 244 | 14.7% | 114 | 14.8% | 174 | 11.3% |
| 61 and older | 459 | 7.4% | 82 | 4.9% | 60 | 7.8% | 96 | 6.2% |
| Male | 2,814 | 45.5% | 711 | 42.9% | 344 | 44.6% | 718 | 46.6% |
| HIV seropositive | 21 | 0.3% | 4 | 0.2% | 5 | 0.7% | 5 | 0.3% |
| Diabetes | 110 | 1.8% | 30 | 1.8% | 16 | 2.1% | 27 | 1.8% |
| BCG scars |  |  |  |  |  |  |  |  |
| 0 | 866 | 14.0% | 211 | 12.7% | 121 | 15.7% | 217 | 14.1% |
| 1 | 3,994 | 64.5% | 1078 | 65.0% | 455 | 59.0% | 974 | 63.2% |
| 2 | 1,053 | 17.0% | 297 | 17.9% | 157 | 20.4% | 283 | 18.4% |
| ≥3 | 276 | 4.5% | 72 | 4.3% | 38 | 4.9% | 67 | 4.3% |
| Smoking status |  |  |  |  |  |  |  |  |
| Non-smoker | 5,745 | 93.8% | 1528 | 93.1% | 724 | 95.0% | 1,443 | 94.3% |
| 1 cigarette per day | 196 | 3.2% | 61 | 3.7% | 22 | 2.9% | 46 | 3.0% |
| >1 cigarette per day | 183 | 3.0% | 52 | 3.2% | 16 | 2.1% | 41 | 2.7% |
| Alcohol use |  |  |  |  |  |  |  |  |
| Non-drinker | 4,479 | 73.8% | 1195 | 73.0% | 562 | 74.5% | 1139 | 75.8% |
| 0 to <3 drinks per day | 1,270 | 20.9% | 353 | 21.6% | 145 | 19.2% | 283 | 18.8% |
| ≥3 drinks per day | 321 | 5.3% | 89 | 5.4% | 47 | 6.2% | 81 | 5.4% |
| Nutritional status |  |  |  |  |  |  |  |  |
| Normal weight | 3,561 | 58.1% | 935 | 57.0% | 422 | 55.2% | 881 | 57.7% |
| Underweight | 103 | 1.7% | 22 | 1.3% | 16 | 2.1% | 27 | 1.8% |
| Overweight | 2,470 | 40.3% | 683 | 41.6% | 327 | 42.7% | 620 | 40.6% |
| Employed | 3,931 | 63.5% | 1019 | 61.5% | 460 | 59.7% | 973 | 63.1% |
| Public transportation | 2,255 | 36.5% | 638 | 38.5% | 311 | 40.3% | 568 | 36.9% |
| Non-user | 1,636 | 26.9% | 469 | 28.8% | 181 | 24.0% | 443 | 29.4% |
| 1 to 3 days per week | 1,969 | 32.4% | 517 | 31.8% | 246 | 32.6% | 495 | 32.9% |
| 4 to 7 days per week | 2,466 | 40.6% | 640 | 39.4% | 328 | 43.4% | 568 | 37.7% |
| Completed high school | 1,143 | 20.7% | 315 | 21.2% | 175 | 25.0% | 247 | 17.8% |
| Socioeconomic status |  |  |  |  |  |  |  |  |
| Low | 2,125 | 35·1% | 602 | 36·9% | 261 | 36·0% | 528 | 34·5% |
| Middle | 2,670 | 44·1% | 682 | 41·8% | 325 | 44·9% | 728 | 47·5% |
| High | 1,261 | 20·8% | 347 | 21·3% | 138 | 19·1% | 276 | 18·0% |

### TB infection

At enrollment, 4,488 (44⋅2%) HHCs were TB-infected. Compared to HHCs of DS-TB index patients, those exposed to INH mono-resistant TB had a 15% (95% CI: 5%-16%) higher risk of infection by 12 months, while those exposed to MDR-TB had an 8% (95% CI: 4-13%) higher risk (Table 2). In sensitivity analyses, the positive association between exposure to both INH mono-resistant TB and MDR-TB and the risk of infection remained similar when we assessed the prevalence ratio of infection at baseline (Table S2), the hazard of TST conversion during follow up (Table S3), when we restricted all of the above analyses to children, and when we estimated the direct effect of DRP on infection by controlling for smear status, cavitary disease, treatment delay, and time to effective treatment (Tables S4-S6).

**Table 2.**
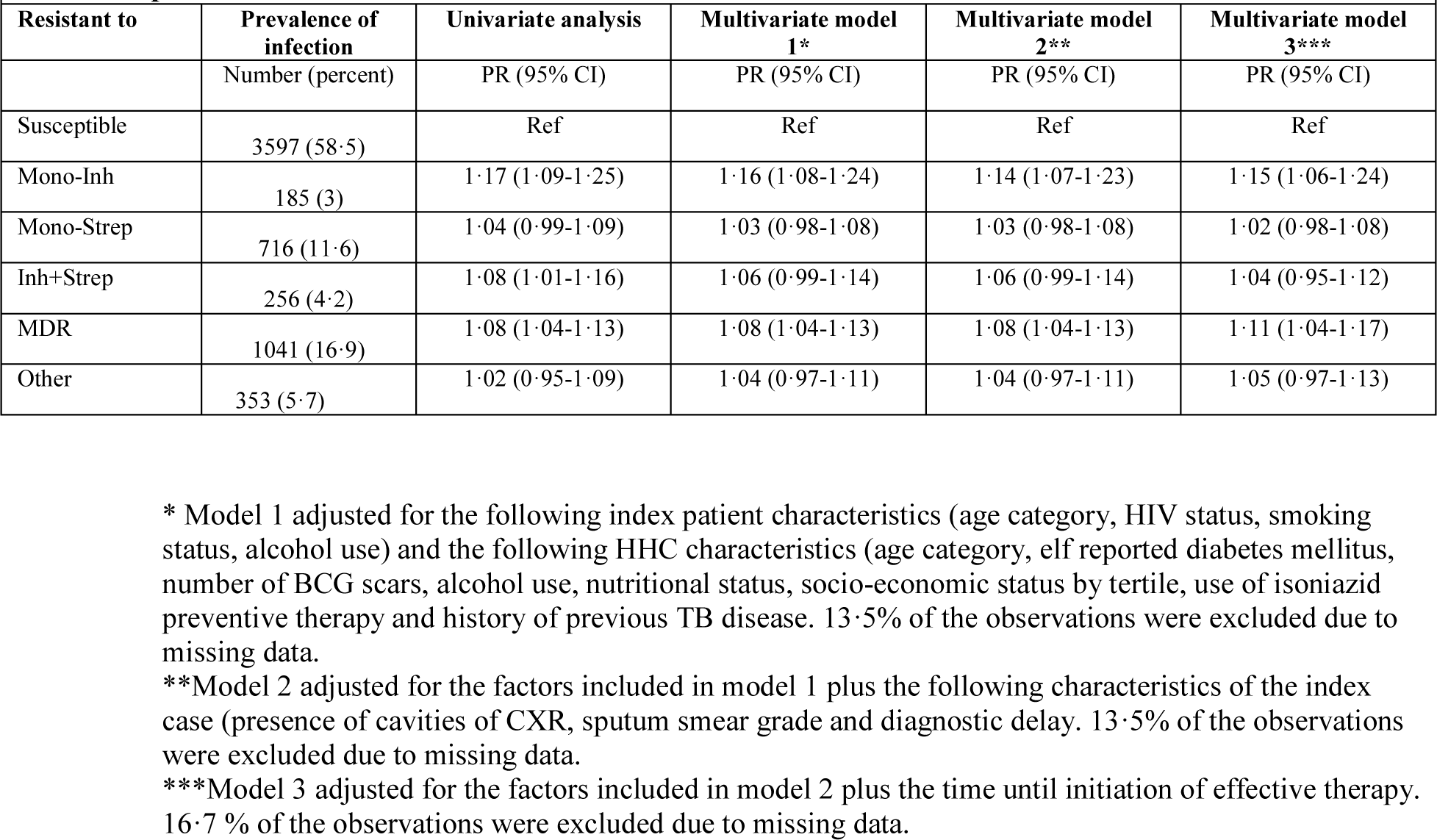
Risk of TB infection among household contacts of TB patients by Mycobacterium tuberculosis drug resistance profile.*

### TB disease

There was no statistically significant difference in the relative hazard of incident TB disease among HHCs exposed to DR-TB compared to DS-TB (Adjusted HR for INH mono-resistant ⋅17 [95% CI: ⋅02-1⋅26]) and for MDR-TB 1⋅28,[(⋅9-1⋅83] (Table 3 and Figure S1). For MDR-TB, this result persisted when we only considered secondary cases if the molecular fingerprint matched that of the corresponding index patient and when matches were based on either more or less stringent criteria (Table 4).

**Table 3.**
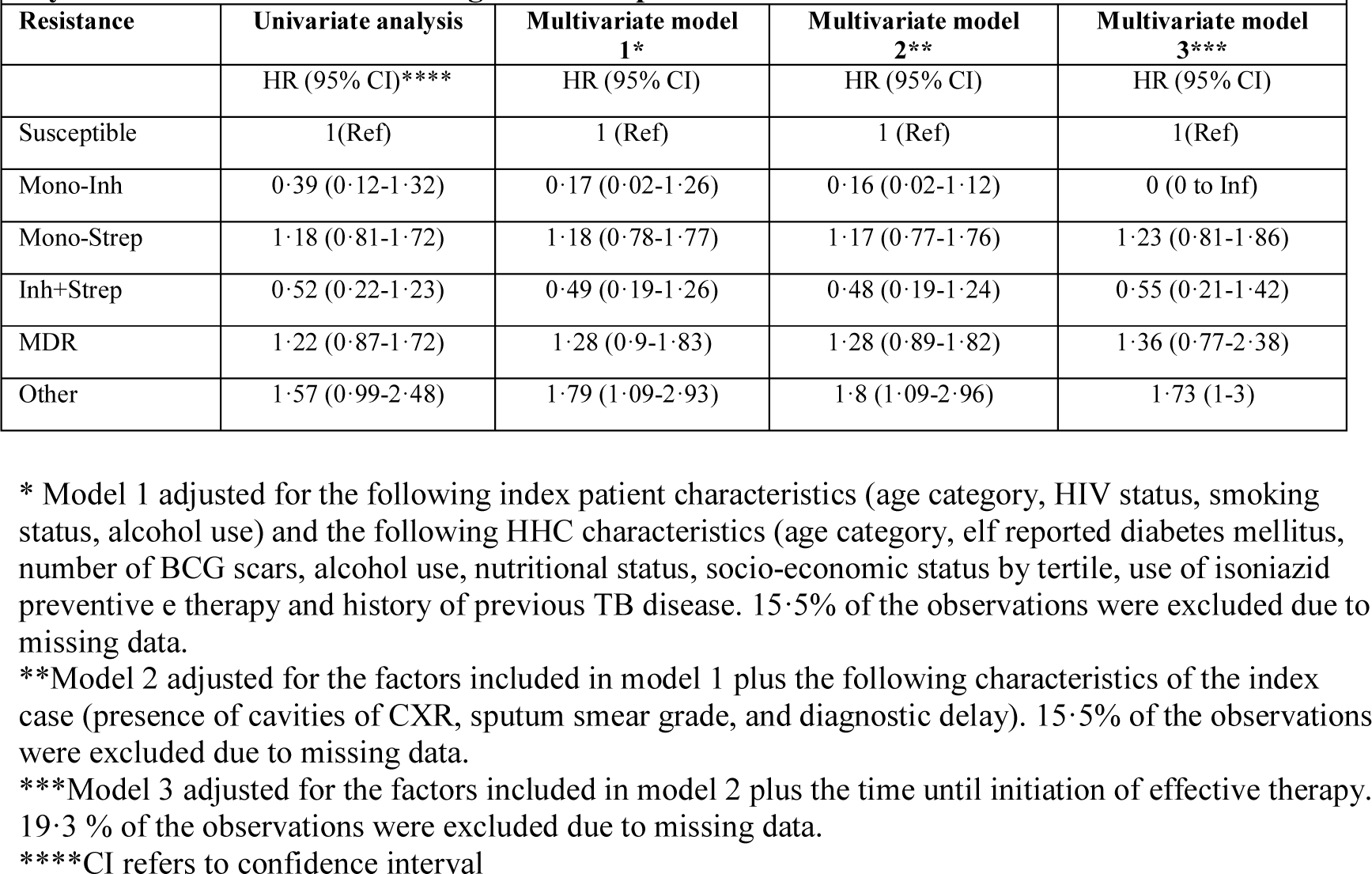
Risk of incident TB disease among household contacts of TB patients by Mycobacterium tuberculosis drug resistance profile.

**Table 4.** Risk of genotypically matched incident TB disease among household contacts of TB patients by Mycobacterium tuberculosis drug resistance profile.

| **Resistance** | **HR (95% CI) \***<br>**(N=10099)** | **HR (95% CI) \*\***<br>**(N=10109)** |
| --- | --- | --- |
| Susceptible | Ref | Ref |
| Mono-Inh | 0 (0-Inf) | 0 (0-Inf) |
| Mono-Strep | 0.53 (0.16-1.77) | 0.45 (0.14-1.51) |
| Inh+Strep | 0.43 (0.05-3.5) | 0.38 (0.05-2.99) |
| Multidrug | 1.13 (0.51-2.5) | 0.98 (0.45-2.12) |
| Other | 1.26 (0.41-3.93) | 1.66 (0.65-4.27) |

\*Hazard ratio for having incident TB that is an identical match with the index patient's MIRU patterns
\*\* Hazard ratio for having incident TB that is a relaxed match with the index patient's MIRU patterns

### TB clustering

Among 3,608 Mtb isolates from unique individuals, 74% were clustered by MIRU genotyping and 26% did not match any other genotype in the dataset (Figure S2). Table 5 shows the relative risk of clustering by potential risk factors for transmission; in both univariate and adjusted analyses, patients with either INH mono-resistant TB or MDR-TB are more likely to be part of a cluster than are DS-TB patients.

**Table 5.** Risk of clustering of MTB genotypes from patients with TB disease by Mycobacterium tuberculosis drug resistance profile.

|  |  |  |  |
| --- | --- | --- | --- |
| **Resistance** | **Univariate**<br>**(N=3608)** | **Multivariate model 1\***<br>**(N=3432)** | **Multivariate model 2\*\***<br>**(N= 2932)** |
|  | RR 95% CI*** | RR 95% CI | RR 95% CI |
| Susceptible | 1 (Ref) | 1 (Ref) | 1 (Ref) |
| Mono-Inh | 1 (0.73-1.37) | 1 (0.73-1.37) | 0.99 (0.72-1.37) |
| Mono-Strep | 0.99 (0.86-1.14) | 0.98 (0.85-1.14) | 1 (0.86-1.16) |
| Inh+Strep | 1.08 (0.9-1.29) | 1.13 (0.95-1.35) | 1.11 (0.92-1.34) |
| Multidrug | 1.27 (1.15-1.42) | 1.24 (1.12-1.38) | 1.18 (0.98-1.42) |
| Other | 1.02 (0.81-1.29) | 1.05 (0.83-1.32) | 1.04 (0.82-1.33) |

\*Model 1 adjusted for the following patient characteristics: gender, HIV status, socioeconomic status by tertile, lineage.
\*\*Model 2 adjusted for the factors in Model 1 plus time to effective treatment.
\*\*\*CI denotes confidence interval

## DISCUSSION

In this large prospective cohort study of TB patient households, we observed that individuals exposed to MDR-TB patients are at higher risk of becoming infected with TB when compared to those exposed to DS-TB and at similar risk of developing TB disease during the 12 months following the diagnosis of the index patient. These data suggest that, compared to DS-TB, MDR-TB is no less transmissible and no less likely to lead to disease. We also found that DR-TB patients are more likely than those with DS-TB to be part of a cluster as defined by a matching MIRU-based genotype. Because clustered patients are presumed to be linked through recent transmission followed by disease progression,^11^ these results support the conclusion that drug resistance does not incur fitness costs in the DR-TB strains circulating in this setting. This hypothesis is also consistent with our observation that the proportion of TB patients who were MDR was highest among the youngest groups, i.e., those most likely to be recently infected.

As the largest study to date of the relative transmissibility and risk of disease progression of DR-TB, our study adds to a diverse body of work on the fitness cost of *Mtb* drug resistance. Laboratory studies that have assessed the fitness cost of drug resistance by comparing bacterial growth rates in media and/or bacillary loads and survival in infected animal models indicate that,^12–14^ while some resistance-causing mutations reduce growth rates or virulence,^15–17^ others have little or variable impact.^18–20^ Even when mutations do confer fitness costs, subsequent “compensatory” mutations can reverse these growth defects while preserving the resistance phenotypes.^21,22^ As would be expected, such low-cost or compensatory mutations are observed at higher frequency than other resistance mutations.^23,24^

Studies of the relative fitness of DR-TB in human populations have approached the question in two ways: by either assessing the effect of DRP on the relative frequency of clustered compared to unclustered cases or by directly measuring the risk of infection and disease among contacts of DS-TB and DR-TB patients.^5^ Multiple molecular epidemiologic studies show that DR-TB strains can be transmitted and that clusters of resistant strains can persist over long periods.^25–28^ However, previous molecular epidemiologic studies have reached different conclusions; some have found that DR strains are more likely to be clustered than DS strains while others have shown the reverse.^29–31^ Some studies have suggested that the association with clustering depends on the specific drug resistance phenotype and/or mutation.^32,33^ Many of these previous studies have been small and subject to biases due to convenience sampling of isolates.^34,35^ Our study, which aimed to prospectively capture all notified cases in a geographically contiguous area, followed by close follow up of household contacts for infection and incidence TB disease, is, to our knowledge, the largest study to systematically examine drug resistance as a risk factor for transmission and disease.

The few studies to date that have directly measured the capacity of DR and DS *Mtb* strains to cause infection or disease have also yielded conflicting results. In 1984, Snider et al. reported no difference in the risk of TB infection in child household contacts exposed to either INH- or streptomycin-resistant *Mtb* compared to DS *Mtb*.^36^ Although the authors also reported no differences in disease incidence in these groups, the study was not powered to detect this outcome and confidence intervals were wide. Similarly, in a small 2001 study conducted in Brazil, Teixeira et al. found that household contacts of MDR-TB patients were slightly more likely to be infected at baseline than contacts of DS-TB patients and equally likely to develop disease, although neither result was statistically significant.^37^ In 2011, the TB Research Centre in India reported on 5,562 household contacts exposed to DS-TB and 779 exposed to INH-resistant TB. This cohort was followed for up to 15 years; the prevalence of TB infection was higher in the contacts of INH-resistant patients, while the hazard of disease was similar in the two groups.^38^ In contrast to these results, in a study also conducted in Peru, Grandjean et al. found that MDR-TB household contacts were half as likely to develop TB disease compared to DS-TB contacts.^39^ Finally, in a small study from Vietnam, Fox et al. found greater TB infection and disease among contacts of patients with known MDR-TB compared to contacts of patients with recently diagnosed TB presumed to be DS-TB (DST was not available for the second group).^40^ Thus, our results are consistent with the majority of previous studies on risk among household contacts, as well as with our own cluster analyses.

We note several important limitations to this study. First, because TSTs only measure previous infection but not the time of its occurrence, there is no ideal way to measure the incidence of TB infection due to a specific, time-dependent exposure. Baseline TST positivity may reflect a remote infection that occurred prior to a person’s exposure to the index patient, while TST conversion among people who tested negative at baseline is subject to survival bias (see Supplement). We examined these issues by conducting multiple sensitivity analyses; all were consistent with our main results.

Secondly, although the study was designed to use molecular fingerprinting to ascertain whether transmission had taken place between an index patient and secondary case, most of the secondary cases in our cohort occurred among child contacts, most of whom did not have microbiological confirmation of TB disease. Although it is expected that a child sick with TB was most likely infected by someone in the home, it is possible that the infection resulted from community-based transmission. Within the subset of 133 secondary cases for whom a genotype was available, only 56 (43⋅6%) matched the genotype of the index patient. Our finding of no significant difference in the hazard of disease after exposure to a DR or DS patient in this subset may reflect the small numbers rather than the absence of an effect. Interestingly, our finding that fewer than half of the genotyped household pairs shared a genotype is consistent with what has been reported from household contact studies in other high burden settings, where this fraction ranged from 25 to 50%.^38,40^ It is unclear whether the incidence of unmatched secondary cases represents background rates of community transmission or signals particularly high levels of vulnerability to TB as a result of shared genetic or environmental risk factors.

These results have major implications for public health policy and for projecting the burden of DR-TB. Mathematical models suggest that the expected trajectories of DS-TB and DR-TB strongly depend on the fitness cost of clinically relevant resistance mutations. If *Mtb* drug resistance exacts no fitness cost, the incidence of DR and MDR TB will be expected to fall more slowly than would be expected even in populations where the acquisition of new drug resistance is minimized through measures such as supervised therapy to ensure adherence to standardized empirical regimens.^41^ Our findings provide strong evidence that TB programs must immediately deploy strategies that directly target DR-TB and MDR-TB, such as the early detection and effective treatment of both infection and disease. These should include both the wider deployment of existing tools and the further development of diagnostic and therapeutic strategies designed specifically for persons already infected with DR-TB.

## Supporting information

## AUTHOR CONTRIBUTIONS

Lima Cohort Study: MBM, MCB, TC, CM and PEF conceived of the study and MBM and MCB led the study design. LL oversaw data collection and management with JB, JTG, RC, ZZ, CC, SA and RY. RC managed laboratory efforts. MBM supervised data analysis and interpretation in conjunction with MCB and CCH.

MCB and MBM wrote the first draft of the manuscript, and all authors contributed to manuscript revision.

## DECLARATION OF INTERESTS

We declare no competing interests.

## ACKNOWLEDGEMENTS

This study was funded by National Institutes of Health (NIH/NIAID CETR U19AI109755).

