## Supplementary material for "Resistance at No Cost: The Transmissibility and Potential for Disease Progression of Drug-Resistant *M. Tuberculosis*"

Supplemental materials

**Table of Contents**

List of Investigators …………………………………………………………….… 2

Methods ………………………………………….………….……............... 3

Results ………………………………………………………………………. 10

Figures ………………………………………………………………………. 10

Tables ……………………………………………………………………….. 11

References ……………………………………..…………………… 18

Appendix ……………………………………………………………………...19

List of Investigators


1. Mercedes C. Becerra, Sc.D., Harvard Medical School, Department of Global Health and Social Medicine, Boston, Massachusetts, United States
2. Chuan-Chin Huang, Sc.D., Brigham and Women's Hospital, Division of Global Health Equity, Department of Medicine, Boston, Massachusetts, United States
3. Leonid Lecca, M.D., Socios En Salud, Lima, Peru
4. Jaime Bayona, M.D., World Bank, Washington, District of Columbia, United States
5. Carmen Contreras, M.P.H., Socios En Salud, Lima, Peru
6. Roger Calderon, M.S., Socios En Salud, Lima, Peru
7. Rosa Yataco, B.S., Socios En Salud, Lima, Peru
8. Jerome Galea, Ph.D., Harvard Medical School, Department of Global Health and Social Medicine, Boston, Massachusetts, United States
9. Zibiao Zhang, M.S., Brigham and Women's Hospital, Division of Global Health Equity, Boston, Massachusetts, United States
10. Sidney Atwood, B.A., Brigham and Women's Hospital, Division of Global Health Equity, Boston, Massachusetts, United States
11. Ted Cohen, D.P.H., Yale School of Public Health, Epidemiology of Microbial Diseases, New Haven, Connecticut, United States
12. Carole D. Mitnick, Sc.D., Harvard Medical School, Department of Global Health and Social Medicine, Boston, Massachusetts, United States
13. Paul Farmer, M.D., Harvard Medical School, Department of Global Health and Social Medicine, Boston, Massachusetts, United States and Brigham and Women's Hospital, Division of Global Health Equity, Boston, Massachusetts, United States
14. Megan Murray, M.D., Harvard Medical School, Department of Global Health and Social Medicine, Boston, Massachusetts, United States and Brigham and Women's Hospital, Division of Global Health Equity, Boston, Massachusetts, United States

### Methods

*Recruitment*

This study was conducted in 106 district health centers in Lima, which are marked on the map in Figure S1. These district-based clinics provide routine health care to people living in their catchments areas and are responsible for most diagnosis and treatment of TB cases. Patients are diagnosed with pulmonary TB disease by health center clinicians on the basis of the Peru national TB guidelines which specify that individuals receive a diagnosis of pulmonary tuberculosis if at least one of two sputum smears is positive for acid-fast bacilli by Ziehl-Neelsen staining, or if a chest radiograph was consistent with tuberculosis in the absence of positive sputum smear results.

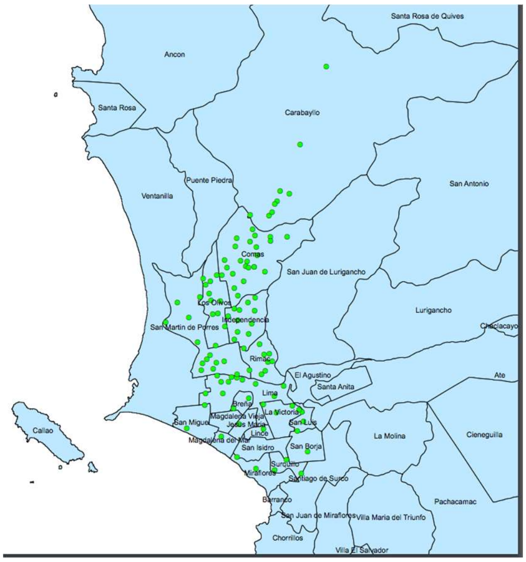


**Figure S1. Study district health centers**

*Enrollment of index patients*

On the day that a health center identified a patient with pulmonary TB, he or she was invited to participate in the study if he/she were over 16 or over and had no cognitive deficits that might interfere with giving informed consent. Once the patient consented to participate, an additional sputum sample was collected and sent for repeat sputum smear microscopy, mycobacterial culture and drug sensitivity testing as described below. We then requested permission to visit the patient's household and recruit his or her household contacts into the prospective cohort study. Study workers aimed to visit households to enroll all household members within one week of the diagnosis of the index case. The median time between index patient's diagnosis and HHC enrollment was 8 days and the interquartile range was 11 days. All individuals who lived in the same household as an index patient at the time the index patient was enrolled in the study were invited to participate. Potential participants were only excluded if they were cognitively unable to provide informed consent or assent.

*Baseline assessment of index patients*

At the time of the index case enrollment, study workers collected the following data through a structured questionnaire: age, gender, occupation, symptoms of tuberculosis, duration of symptoms, history of previous TB disease, alcohol, intravenous drug and tobacco history, co-morbidities including HIV status and DM. Patients who did not know their HIV status had blood drawn for HIV and CD4 count. Signs associated with TB disease, height and weight were recorded.

*Ascertainment of TB disease*

*Bacteriological cultures*

The study staff noted results of routine sputum smear microscopy at the participating health centers. Health centers also sent sputum samples for a minority of index patients who had known risk factors for drug-resistant TB for routine culture and DST at the designated public health laboratory for each center. In addition to this routine diagnostic microbiology, a sputum sample was sent to the designated research laboratory (Lima Ciudad Regional Reference Lab: September 2009 – November 2010; Blufstein Laboratorio Clínico from November 2010 to March 2012, Socios en Salud Laboratory from March to August 2012) for repeat smear, culture and drug sensitivity testing.

Sputum samples were transported in containers kept at 2-8°C and bacteriological examination was carried out as soon as possible. Samples were decontaminated and further homogenized, following the NALC method, centrifuged at 2000*g* to 3000*g* for less than 20 minutes. The resulting sediments were then neutralized in phosphate buffer solution and re-suspended. Sediments were tested for the presence of acid-fast bacilli by Ziehl-Neelsen staining and cultured by inoculation into 2 tubes containing Lowenstein-Jensen or Ogawa medium. Between March to August 2012, some specimens were also cultured using the MODS assay to allow more rapid identification of microbiologically confirmed cases. Inoculated tubes were incubated at 37°C until bacterial growth (e.g., colonies on solid media or cord formation in MODS medium) for 60 days or until colonies were found, whichever came first. Isolates in which we observed any possible discrepancy with MTB complex, were assessed using biochemical identification tests. After culture, all colonies from isolates were harvested on cryovials with 7H9 Middlebrook broth with 20% of glycerol. Those were then incubated overnight at 4 Celsius degrees and stored at -60 to -80°C.

*Drug-susceptibility testing*

Indirect susceptibility testing to INH, Rifampicin, Ethambutol and Streptomycin was conducted by the Löwenstein-Jensen proportion method, using the following drug concentrations: isoniazid (0·2 and 1·0 µg/ml), rifampin (40·0 µg/ml), ethambutol (2.0 µg/ml), and streptomycin (4·0 µg/ml) and to pyrazinamide PZA (100 ug/ml) by the Wayne method. For those isolates in which any resistance to first line drugs was detected, susceptibility testing to the following panel of second-line drugs was also performed by the agar plate method, using the following drug concentrations: kanamycin (6·0 µg/mL), capreomycin (10·0 µg/mL), ethionamide (10·0 µg/mL), ciprofloxacin (2·0 µg/mL), para-aminosalicylic acid (8·0 µg/mL), cycloserine (30·0 µg/mL).

Drug-susceptibility test results were recorded in a secure web-based information management system that is used by the laboratory and paper results were sent to the study data center for distribution to the health center, the participant, and the participant´s study file.

*Quality assurance*

To ensure that the results of susceptibility testing were reliable, a system of quality assurance was implemented. As an internal quality control, a standard H37Ra strain of *M. tuberculosis* was tested against the entire drug panel with each new batch of media.

External quality control for drug-susceptibility testing was done in conjunction with the National Mycobacteria Reference Laboratory (National Institute Of Health in Peru) and The College of American Pathologist which provided panel of coded strains for blind retesting for ZN smear; culture, MTB identification and drugs susceptibility testing for first and second line drugs. Results of yearly blind testing are presented in Appendix 1.

*TB DNA fingerprinting*
DNA isolates were extracted from mycobacterial colonies grown in culture media (Lowenstein-Jensen, Ogawa, or MODS) and transported from Peru to Genoscreen, France. Upon receipt, all specimens were entered into the study database by scanning of the bar-codes on the vials. The specimens were stored between -60°C to -80°C.

*DNA isolation and molecular genotyping.*
DNA was extracted and genotyped by 24-loci mycobacterial interspersed repetitive units–variable-number tandem repeats (MIRU-VNTR) using standard methods.[1]

*Chest radiograph*y
Chest radiographs were performed at health center or at local imaging facilities. Films were digitized in the study database, with the subject's identifying information removed, and labeled with the unique study subject identification number. Two radiologists read each film and completed a standardized form. Chest films were then given to the health center to be filed in the subject's medical chart.

*Ascertainment of HIV Infection*
Peru NTP guidelines specify that all TB patients undergo HIV testing. From September 2009 to August 2011, HIV infection status was determined using a lab-based enzyme immunoassay (EIA), and nonnegative samples were confirmed using an immunofluorescence assay (IFA). After August 2011, a new HIV testing algorithm was employed;[2] we first performed a rapid screening test and followed those with nonnegative tests with an EIA and a confirmatory IFA. Study staff coordinated the appropriate confirmatory tests for participants whose results were "reactive" or "indeterminate." All study participants had pre-HIV counseling by trained study staff or Ministry of Health personnel prior to the collection of a blood specimen as well as post-HIV test counseling after testing, according to the guidelines of the National STI/HIV/AIDS Program. Participants were informed of their results by a trained study worker or a trained staff member at the Ministry of Health clinic.

*Follow up of Index patients*
Index patients received directly observed therapy at their district health clinics as specified in the Peru NTP guidelines for drug sensitive and drug resistant TB. Patients with drug sensitive TB received a standard 6 month course with a 2 month "intensification phase" of INH, Rifampicin, PZA and Ethambutol followed by a four month "consolidation phase" of INH and Rifampicin alone. Patients with MDR TB received treatment according to Peru NTP guidelines. Since results for routine drug resistance testing were often not available for 2-3 months after initial diagnosis, patients who were not previously suspected of having MDR TB were started on a first line drug regimen until the diagnosis of MDR was confirmed. Thus, many patients with DR TB did not start "effective therapy" until several months into their treatment course. (see definition of effective therapy below under Analyses). Study staff collected follow-up data for those with DS TB at 2, 6, 12 and 24 months and for those with DR TB again at 36 or 48

months. This included a record of all drug treatment including any changes in drug regimens and the dates when these occurred.

At two and six months, patients were evaluated by repeat sputum smear microscopy and culture. Culture positive sputum samples underwent repeat drug sensitivity testing and DNA genotyping as described above.

*Enrollment of household contacts*
At the time of the enrollment of household contacts, study workers collected the following data through a structured questionnaire: age, gender, relationship to index case, housing information including number of rooms, building material, type of flooring, income, education, history of incarceration, occupation, alcohol, cigarette and illicit drug intake, general health history including previous history of TB, BCG vaccination, co-morbidities including HIV and DM, and medications taken. Participants were queried about the presence of any symptoms associated with TB disease including cough, night sweats, weight loss or fever. Participants who reported any of these symptoms were referred their local health clinic for chest radiography and clinical evaluation for active TB disease. They were also asked if they had been offered and initiated isoniazid preventive therapy.

Trained study staff measured height and weight of all participants. All household members received a tuberculin skin test, with the exception of: (1) subjects with active TB disease or a history of TB disease, (2) subjects who may have severe eczema, (3) subjects known to be hypersensitive to any component of the Tuberculin PPD RT23 SSI (i.e., those who tested positive in the past) and (4) subjects who previously have experienced an adverse reaction to tuberculin products. Tests were placed intradermally on the forearm and were read between 48 and 72 hours after intradermal injection. The diameter of induration was measured transversely to the long axis of the forearm and recorded in millimeters.

Participants were offered HIV pretest counseling and testing. Prior to August 2011, blood samples were collected at the contacts' homes and sent to a laboratory for enzyme-linked immunosorbent assay testing. Negative results were delivered to the HHCs 2–4 weeks later, while participants with non-negative results were linked to a Ministry of Health clinic for confirmatory sample extraction and follow-up. From August 2011 on, study staff performed home based HIV testing using the Determine® HIV 1/2 Ag/Ac Combo test* (Alere, Jouy-en-Josas, France).

*Follow-up of household contacts*
Participants were asked to advise study staff if they were diagnosed with active TB disease prior to the next scheduled study follow up visit. Participants were re-visited in their household at 2, 6 and 12 months at which times they were asked about any TB diagnoses that had occurred as well as about any symptoms of active disease. Those who reported symptoms were referred to their local health center for further clinical evaluation including a chest radiograph and sputum smear. Participants who tested negative at the initial study visit and who had not developed active TB disease at the time of the follow up visit were underwent repeat TST at 6 and 12 months.

Analyses
The time to treatment was measured as the number of days the patient reported coughing prior to diagnosis. Time to effective therapy was measured as the time from

diagnosis of the patient until he or she received a drug regimen deemed appropriate for his or her DR. For patients with MDR or XDR TB, we considered a drug regimen "effective" if it included at least four drugs to which the patient's isolate was susceptible. If a strain was resistant to at least one drug other than Rifampicin, we considered a regimen to be effective if it included at least three effective drugs, one of which was Rifampicin. If a strain was resistant to Rifampicin but not INH or PZA, we considered a regimen to be effective if it included at least three drugs to which strain was susceptible, one of which was PZA.

We categorized participants according to their alcohol intake as nondrinkers if they reported having consumed no alcoholic drinks per day), light drinkers if they reported drinking <40 grams or <3 alcoholic drinks per day and heavy drinkers if they reported drinking 40 grams of alcohol or more or 3 or more drinks per day. A large proportion of smokers reported smoking only a single cigarette per day. We classified people as nonsmokers if they reported no cigarette smoking, as light smokers if they reported smoking one cigarette per day and as heavy smokers if they reported smoking more than one cigarette per day. We defined nutritional status for children based on the World Health Organization body mass index (BMI) z-score tables.[3] We assigned people with BMI *z*-scores of less than 2 as underweight and those greater than 2 as overweight.

We created a continuous variable to capture summarize household-level socioeconomic status by including variables on housing quality, water supply and sanitation in a principal component analysis (PCA) (as described in 4.) PCA is a data reduction statistical technique that extracts a set of uncorrelated 'principal components' from a set of correlated variables, where each principal component is a weighted linear combination of the original variables. Appendix 2 provides the variables which were used to generate a composite SES score as a continuous variable and their weight in the first principal component. The continuous SES score was categorized into tertiles corresponding to relative "low," "middle," and "upper" SES.

Outcomes

Infection

We considered the following infection outcomes in our analyses: infection among HHCs at baseline, infection during 12 months of follow up among HHCs who were uninfected at baseline, and infection by 12 months of follow-up. We considered people infected at baseline if they had a history of prior TB disease, reported a history of a positive TST or had a positive TST at baseline. We considered people to have become infected with TB during follow up if they had a negative TST at baseline and a positive TST at some point during follow up.

Disease

We identified incident TB by direct household visits and from medical records from the hospitals within the study area. We considered HHCs to have co-prevalent TB if they were diagnosed within two weeks of the diagnosis of the index case and to be "secondary" cases if they were diagnosed between days 15 and days 455 of follow-up (90 days extensive buffer time for the 12 months visit). Diagnosis of adult secondary TB followed the same criteria as outlined above for index cases. We assumed that we captured all incident cases, so all HHCs without evidence of progressive TB disease were considered disease-free at the end of follow-up. In addition, we defined secondary TB disease among contacts younger than 18 years of age according to the consensus guidelines for classifying TB disease in children.[5]

Data Analysis.

Infection at baseline and the end of follow-up

Because TB is a disease with an insidious onset, it is likely that many patients have had infectious TB for weeks or months prior to their presentation to a health facility for diagnosis. Thus, HHCs are likely to have been exposed to the index case during an infectious period of unknown duration and some of these will have become infected during this exposure prior to the time that the index case presents for diagnosis. Although we would expect that HHCs exposed to index cases who were more likely to transmit TB would have a higher prevalence of recent infection at the time of the diagnosis of the index case than those exposed to less infectious index cases, the tuberculin skin test does not distinguish between recent and more remote infection. An assessment of relative transmissibility of an index case that is based on the prevalence odds of TB infection among the exposed and unexposed at baseline is therefore subject to bias by non-differential misclassification if a positive TST is taken as evidence of a recent transmission event. We have nonetheless chosen to report this estimate in addition to the risk of TST conversion from negative to positive conditional on the exposure because the TST negative population at the time of enrollment constitutes a “survival cohort” that may be depleted of those at highest risk for conversion. Thus, a very strong index patient risk factor for TB transmission might have led to earlier TST conversion, i.e., prior to the diagnosis of disease in the index case, and the population of HHCs with such an exposure who remain uninfected at baseline may include “infection resistors,” people who experience “early clearance” of TB infection and do not mount a cell mediated immune response that leads to TST conversion.[6]

We estimated prevalence ratios for infection at baseline and by the end of follow-up using a modified Poisson generalized estimating equation to account for correlation among participants within a household. We specified an exchangeable working correlation structure for observations within the same household. For inference, we obtained empirical standard error estimates that were used to construct Wald type 95% confidence intervals. We first performed age-adjusted univariate analyses for covariates that were potential predictors of tuberculosis infection based on a priori background knowledge. Subsequently, all the covariates were entered into a backwards stepwise algorithm, with the exception of sputum smear status, length of symptomatic period, presence of cavitary disease, and time to effective treatment of index case, as we hypothesized that these risk factors may mediate the effects of the index patient’s drug resistance status on the risk of tuberculosis infection among HHCs. We retained variables with $p<0{\cdot}1$ and variables which were likely to modify TB infection (HIV status of index case, smoking and drinking status of index case, and socioeconomic status of household) in the multivariate model. We also aimed to quantify the direct effect of DRP on risk of tuberculosis transmission, i.e., the effect which was not mediated by smear status, duration of symptomatic disease, cavitation, or time to effective treatment of index case. We evaluated the direct effect by adding the mediators to the regression model, assuming that upon adjusting for the observed covariates, no unobserved confounding prevailed for the joint effects of the degree of the index patient’s immunosuppression and these 4 mediators on the HHC’s risk of tuberculosis infection. As noted above, this analytic approach implicitly assumes that all infected HHCs acquired infection from the index patient. If some HHCs acquired infection elsewhere, this would lead to non-differential misclassification of the exposure status and would be expected to attenuate our results toward the null. To address this, we conducted a sensitivity analysis restricting the analyzed cohort to children who we assumed were less likely to be infected in the past or in the community than adults.

2. Time to infection

We measured time from enrollment to infection among HHCs who were uninfected at baseline. We defined the date of infection as the midpoint between date of enrollment and the date of a positive TST result and we censored contacts who remained TST negative at the date of last TST result. We used a Cox frailty proportional hazards models to evaluate risk factors for incident TB infection, accounting for clustering within households. In the multivariate model, we included variables identified a priori as potential confounders (HIV status of index case, smoking and drinking status of index case, and socioeconomic status of household) and any others associated with the outcome with a p value < 0·1 using a backwards stepwise algorithm. We verified the proportional hazards assumptions for each covariate by introducing an interaction term between the covariate and time; we stratified by variables for which the proportional hazards assumption did not hold. We followed the same logic described above evaluating the mediation effects of the four possible mediators, smear status, duration of symptomatic disease prior to diagnosis, cavitation on chest film, or time to effective treatment of index case. We repeated the sensitivity analysis described above restricting our analysis to child HHC based on the assumption that child contacts are more likely to be infected within household than adults.

Disease incidence

We measured the time from enrollment to disease occurrence among all HHCs except those with co-prevalent TB. We used a Kaplan-Meier curve to examine the disease-free survival time and Cox frailty proportional hazards models to evaluate risk factors for incident TB disease, accounting for clustering within households. In the multivariate model, we included variables identified a priori as potential confounders (HIV status of index case, smoking and drinking status of index case, socioeconomic status of household, and IPT) and any others associated with the outcome with a p value < 0·1 using a backwards stepwise algorithm. We followed the same logic described above evaluating the mediation effects of the four mediators. In a sensitivity analysis, we considered only those secondary patients who shared a MIRU pattern with the index case, excluding from the analysis those with secondary disease who did not share an index case. We conducted this sensitivity analysis in two different ways. In the first, we considered secondary cases whose 24 locus MIRU patterns were an exact match with the index case and in the second, we considered secondary cases whose MIRU patterns matched on 22 of 24 loci. Because the number of incident TB cases was low in these sensitivity analyses, we only performed the univariate analyses.

Cluster analysis

We genotyped the sputum samples of the microbiologically confirmed cases (including index, co-prevalent, and secondary cases) using the 24-locus-MIRU-VNTR. Some cases had more than one sputum sample because their sputum samples were collected multiple times at different time points. We excluded the samples if they had mixed alleles in any loci or missing MIRU typing on more than one locus. Samples sharing identical 24-MIRU typing with at least one other sputum sample collected from a different case were considered as “identical match clustered” samples, assuming missing alleles could match with any allele. Samples with unique MIRU typing were considered “non-

clustered”. In the sensitivity analyses, we used a relaxed matching process, allowing up to 2 mismatched MIRU loci in defining the “relaxed match clustered” samples.
We examined whether the samples’ drug resistance profiles were associated with being “clustered” or not using a modified Poisson generalized estimating equation to account for the repeated sputum sample collections. We first performed univariate analyses for covariates that were potential predictors of clustering. In the multivariate model, we included variables identified a priori as potential predictors (age, gender, HIV status, socioeconomic status of household) and any other variables associated with the outcome with a p value < 0·1 using a backwards stepwise algorithm. We did not adjust for time to effective therapy in the multivariate model because we did not collect this variable’s information for the secondary or co-prevalent cases.

**Results**
In addition to the prevalence of infection by 12 months reported in the main text, we also assessed the prevalence odds of infection at baseline by drug resistance profile both in the entire household contact cohort and in the subset of children younger than fifteen years Table S2 and S3 show that in both the entire cohort and among children, contacts exposed to patients with INH mono-resistant or MDR TB were more likely to be infected at baseline. Other resistance phenotypes had no impact on the prevalence of infection at baseline. Point estimates for mono-INH and MDR were further amplified in the subset of the cohort aged 15 and under.

We also assessed the hazard ratio of TST conversion among individuals who were TST negative at baseline. This analysis confirmed our finding that contacts of MDR index patients are at higher risk of being infected in the entire cohort (Table S4) and among children (Table S5), but showed no difference between rates of conversion among contacts exposed to other types of resistance.

**Cluster Distribution**

**Figure S2 shows the distribution of cluster sizes among microbiologically confirmed patients with TB.**

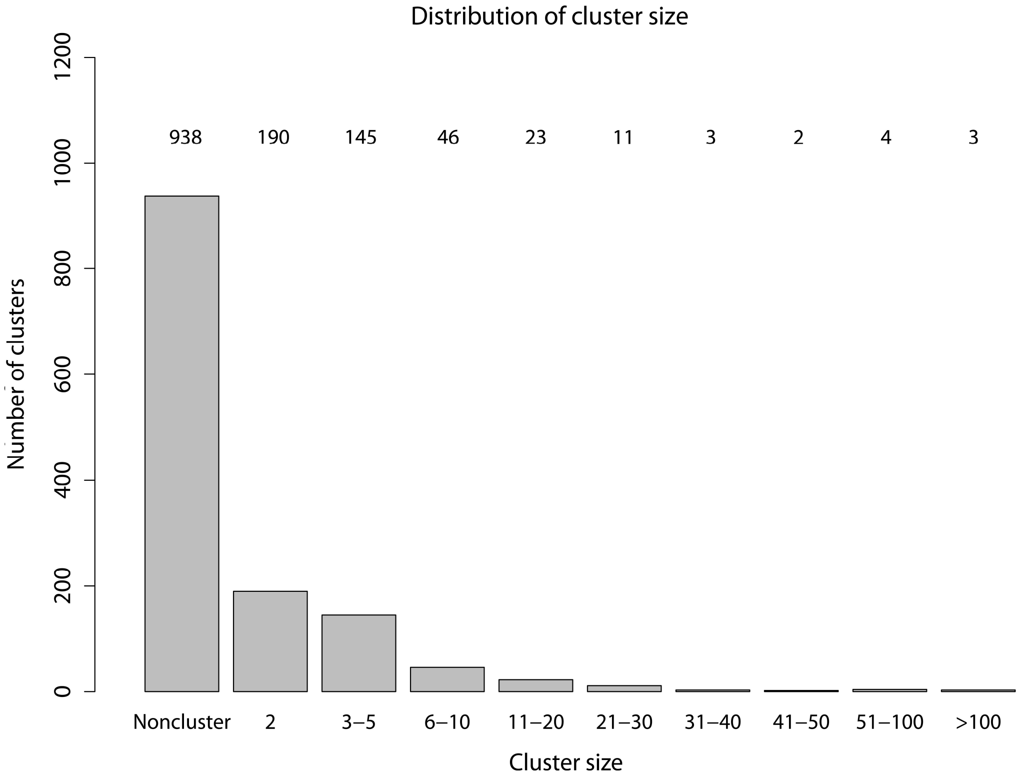

### Tables

| Table S1. **Characteristics of index TB patients in Lima, Peru** | |
|---|---|
| **Characteristic (total *n* with data)** | **N (percent)*** |
| Age, years (n=4,044) | |
| 16 to 30 | 2,262 (55·9) |
| 31 to 45 | 885 (21·9) |
| 46 to 60 | 519 (12·8) |
| 61 and older | 378 (9·3) |
| Male gender (n=4,044) | 2,503 (61·9) |
| HIV seropositive (n=3,986) | 138 (3·5) |
| Sputum smear status (n=4,026) | |
| Negative | 1077 (26·8) |
| + | 1057 (26·3) |
| ++ | 735 (18·2) |
| +++ | 1157 (28·7) |
| Treatment delay (n=3,970) | |
| < 2 weeks | 893 (22·5) |
| 2-4 weeks | 1265 (31·9) |
| 4-8 weeks | 987 (24·9) |
| > 8 weeks | 825 (20·8) |
| Cavitary lesion (n=3,961) | 1082 (27·3) |
| Drug resistance profile (n=3,339) | |
| Susceptible | 2,065 (61·8) |
| Mono-resistant | 538 (16·1) |
| Poly-resistant | 258 (7·7) |
| Multidrug-resistant | 478 (14·3) |
| Drug resistance profile 2 (n=3,339) | |
| Susceptible | 2065 (61·8) |
| Mono-Inh resistant | 85 (2·5) |
| Mono-Strep resistant | 378 (11·3) |
| Inh+Strep resistant | 129 (3·9) |
| Multidrug-resistant | 478 (14·3) |
| Other types of resistance | 204 (6·1) |
| Smoking status (n=3,959) | |

| | |
|---|---|
| Non-smoker | 3,859 (97·5) |
| 1 cigarette per day | 35 (0·9) |
| >1 cigarette per day | 65 (1·6) |
| Alcohol use (n=3,869) | |
| Non-drinker | 2,197 (56·8) |
| 0 to <3 drinks per day | 1270 (32·8) |
| ≥3 drinks per day | 402 (10·4) |

Table S2. **Risk of TB infection at baseline among household contacts of TB patients by MTB drug resistance profile.**

| **Resistant to** | **Prevalence of infection** | **Univariate analysis** | **Multivariate model 1*** | **Multivariate model 2*** | **Multivariate model 3*** |
|---|---|---|---|---|---|
| | Number (percent) | PR (95% CI)**** | PR (95% CI) | PR (95% CI) | PR (95% CI) |
| Susceptible | 2,635 (58·7) | Ref | Ref | Ref | Ref |
| Mono-Inh | 149 (3·3) | 1·41 (1·25-1·59) | 1·4 (1·23-1·59) | 1·37 (1·21-1·56) | 1·47 (1·3-1·66) |
| Mono-Strep | 536 (11·9) | 1·06 (0·98-1·15) | 1·03 (0·95-1·11) | 1·02 (0·94-1·11) | 1·03 (0·95-1·12) |
| Inh+Strep | 189 (4·2) | 1·08 (0·96-1·21) | 1·03 (0·91-1·16) | 1·02 (0·91-1·15) | 0·99 (0·86-1·13) |
| MDR | 729 (16·2) | 1·09 (1·02-1·17) | 1·09 (1·01-1·17) | 1·09 (1·01-1·17) | 1·17 (1·05-1·31) |
| Other | 250 (5·6) | 1·01 (0·9-1·13) | 1·03 (0·92-1·16) | 1·03 (0·92-1·16) | 1·05 (0·93-1·19) |

*Model 1 adjusted for the following characteristics of the index patient (age category, gender, HIV status, smoking status, alcohol use) and of the HHC (age category, BCG scares, smoking status, alcohol use, and SES tertile). 13•4% of the observations were excluded due to missing data.
**Model 2 adjusted for the factors included in model 1 plus the following characteristics of the index case (presence of cavities of CXR, sputum smear grade and diagnostic delay. 13•4% of the observations were excluded due to missing data.
***Model 3 adjusted for the factors included in model 2 plus the time until initiation of effective therapy. 17•3 % of the observations were excluded due to missing data.
****CI refers to confidence interval

Table S3. **The hazard of TB infection among initially uninfected household contacts of TB patients by MTB drug resistance profile.**

| Resistance | Univariate analysis | Multivariate model 1* | Multivariate model 2** | Multivariate model 3*** |
|---|---|---|---|---|
| | HR (95% CI) | HR (95% CI) | HR (95% CI) | HR (95% CI) |
| Susceptible | Ref | Ref | Ref | Ref |
| Mono-Inh | 1·4 (0·92-2·13) | 1·22 (0·70-2·11) | 1·17 (0·67-2·02) | 1·16 (0·67-2·01) |
| Mono-Strep | 1·13 (0·92-1·38) | 1·01 (0·81-1·27) | 1·02 (0·82-1·28) | 1·03 (0·83-1·29) |
| Inh+Strep | 1·09 (0·79-1·49) | 1·05 (0·72-1·54) | 1·03 (0·70-1·50) | 1·03 (0·71-1·52) |
| MDR | 1·42 (1·2-1·68) | 1·44 (1·19-1·75) | 1·44 (1·18-1·74) | 1·58 (1·15-2·18) |
| Other | 1·13 (0·87-1·47) | 1·11 (0·81-1·51) | 1·12 (0·82-1·52) | 1·16 (0·84-1·59) |

* Model 1 adjusted for the following characteristics of the index patient (age category, gender, HIV status, smoking status and alcohol use) and of the household contact (age category, number of BCG scars, smoking status, alcohol use and socioeconomic status tertile).
** Model 2 adjusted for the factors included in Model 1 plus sputum smear status, cavitary lesions, and diagnostic delay.
*** Model 3 adjusted for the factors included in Model 2 and for time to effective treatment.
****CI denotes confidence interval.

Table S4. **Risk of TB infection among child household contacts of TB patients by Mycobacterium tuberculosis drug resistance profile.***

| Resistant to | Prevalence of infection | Univariate analysis | Multivariate model 1 | Multivariate model 2 | Multivariate model 3 |
|---|---|---|---|---|---|
| | Number (percent) | PR (95% CI) | PR (95% CI) | PR (95% CI) | PR (95% CI) |
| Susceptible | 851 (58·4) | Ref | Ref | Ref | Ref |
| Mono-Inh | 43 (2·9) | 1·27 (1·01-1·59) | 1·31 (1·04-1·65) | 1·24 (0·99-1·55) | 1·27 (1·01-1·6) |
| Mono-Strep | 186 (12·8) | 1·09 (0·96-1·25) | 1·09 (0·95-1·24) | 1·08 (0·94-1·23) | 1·1 (0·956-1·26) |
| Inh+Strep | 48 (3·3) | 1·03 (0·83-1·28) | 1·09 (0·87-1·37) | 1·06 (0·85-1·33) | 1·07 (0·85-1·36) |
| MDR | 240 (16·5) | 1·11 (0·99-1·25) | 1·14 (1·01-1·29) | 1·12 (1-1·26) | 1·21 (1-1·46) |
| Other | 89 (6·1) | 1·11 (0·93-1·32) | 1·11 (0·92-1·34) | 1·11 (0·93-1·33) | 1·13 (0·93-1·37) |

* Model 1 adjusted for the following characteristics of the index patient (age category, gender, HIV status, smoking status and alcohol use) and of the household contact (age category, number of BCG scars, smoking status, alcohol use and socioeconomic status tertile).
** Model 2 adjusted for the factors included in Model 1 plus sputum smear status, cavitary lesions, and diagnostic delay.
*** Model 3 adjusted for the factors included in Model 2 and for time-effective treatment.
****CI denotes confidence interval.

Table S5. **Risk of TB infection at baseline among child household contacts of TB patients by MTB drug resistance profile.***

| **Resistant to** | **Prevalence of infection** | **Univariate analysis** | **Multivariate model 1** | **Multivariate model 2** | **Multivariate model 3** |
|---|---|---|---|---|---|
| | Number (percent) | PR (95% CI) | PR (95% CI) | PR (95% CI) | PR (95% CI) |
| Susceptible | 573 (58·7) | Ref | Ref | Ref | Ref |
| Mono-Inh | 34 (3·5) | 1·57 (1·18-2·09) | 1·66 (1·23-2·24) | 1·56 (1·17-2·09) | 1·78 (1·35-2·35) |
| Mono-Strep | 139 (14·2) | 1·23 (1·03-1·46) | 1·17 (0·97-1·4) | 1·16 (0·96-1·39) | 1·18 (0·98-1·43) |
| Inh+Strep | 27 (2·8) | 0·92 (0·64-1·31) | 0·92 (0·63-1·36) | 0·89 (0·61-1·3) | 0·91 (0·61-1·33) |
| MDR | 145 (14·9) | 1·05 (0·88-1·25) | 1·06 (0·89-1·27) | 1·04 (0·87-1·25) | 1·21 (0·88-1·67) |
| Other | 58 (5·9) | 1·1 (0·85-1·42) | 1·06 (0·81-1·39) | 1·05 (0·8-1·37) | 1·06 (0·79-1·42) |

Table S6. **The hazard of TB infection among initially uninfected child household contacts of TB patients by MTB drug resistance profile.**

| **Resistance** | **Univariate analysis** | **Multivariate model 1*** | **Multivariate model 2**** | **Multivariate model 3***** |
|---|---|---|---|---|
| | HR (95% CI)**** | HR (95% CI) | HR (95% CI) | HR (95% CI) |
| Susceptible | Ref | Ref | Ref | Ref |
| Mono-Inh | 1·07 (0·51-2·28) | 0·85 (0·34-2·13) | 0·74 (0·29-1·89) | 0·74 (0·29-1·87) |
| Mono-Strep | 1·06 (0·74-1·52) | 1·13 (0·78-1·64) | 1·13 (0·77-1·64) | 1·15 (0·79-1·68) |
| Inh+Strep | 1·23 (0·72-2·12) | 1·51 (0·82-2·78) | 1·51(0·81-2·78) | 1·51 (0·82-2·78) |
| MDR | 1·52 (1·15-2·01) | 1·57 (1·15-2·15) | 1·56 (1·14-2·14) | 1·77 (1·01-3·08) |
| Other | 1·31 (0·83-2·05) | 1·38 (0·84-2·27) | 1·40 (0·85-2·30) | 1·46 (0·88-2·43) |

* Model 1 adjusted for the following characteristics of the index patient (age category, gender, HIV status, smoking status and alcohol use) and of the household contact (age category, number of BCG scars, smoking status, alcohol use and socioeconomic status tertile).
** Model 2 adjusted for the factors included in Model 1 plus sputum smear status, cavitary lesions, and diagnostic delay.
*** Model 3 adjusted for the factors included in Model 2 and for time to effective treatment.
****CI denotes confidence interval.

**Appendix 1: Annual Quality Control results for blind panel testing of Mtb samples**

**The College of American Pathologist:**

2012- Satisfactory 100%
AFB Quantification: 100%
First line DST: 100%
MTB Identification: 100%
AFB Smear Stain: 100%
Mycobacteria Screen (Culture): 100%

2013-A- Satisfactory 100%
AFB Quantification: 100%
Second line DST: 100%
MTB Identification: 100%
AFB Smear Stain: 100%
Mycobacteria Screen (Culture): 100%

2013-B- Satisfactory 95%
AFB Quantification: 100%
First line DST: 80%
MTB Identification: 100%
AFB Smear Stain: 100%
Mycobacteria Screen (Culture): 100%

2014-A- Satisfactory 100%
AFB Quantification: 100%
Second line DST: 100%
MTB Identification: 100%
AFB Smear Stain: 100%
Mycobacteria Screen (Culture): 100%

2014-B- Satisfactory 100%
AFB Quantification: 100%
First and Second line DST: 100%
MTB Identification: 100%
AFB Smear Stain: 100%
Mycobacteria Screen (Culture): 100%

**National Institute of Heath - PERU**

EQA 2013-2014:

Drug (SENSITIVITY)-(SPECIFICITY)
RIF: 100% - 100%
INH: 100% - 100%
ETB: 85.7% - 94.1%
CIP: 100% - 87.5%
KAN: 100% - 87.5%
CAP: 100% - 94.7%
ETH: 61.5% - 94.1%
